# Epidemiology, ecology and human perceptions of snakebites in a savanna community of northern Ghana

**DOI:** 10.1101/543009

**Authors:** Yahaya Musah, Evans P.K. Ameade, Daniel K. Attuquayefio, Lars H. Holbech

**Affiliations:** Department of Animal Biology and Conservation Science, University of Ghana, P.O. Box LG 67, Legon, Accra, Ghana.; Department of Pharmacology, University for Development Studies, P.O. Box TL 1350, Tamale, Ghana.

**Keywords:** Serpentes, envenomation incidence, demography, Guinea savannah, West Africa

## Abstract

**Background:** Worldwide, snakebite envenomations total ∼2.7 million reported cases annually with ∼100,000 fatalities. Since 2009, snakebite envenomation has been classified as a very important ‘neglected tropical disease’ by the World Health Organisation. Despite this emerging awareness, limited efforts have been geared towards addressing the serious public health implications of snakebites, particularly in sub-Saharan Africa, where baseline epidemiological data remain incomplete. Due to poverty as well as limited infrastructure and public health facilities, people in rural Africa, including Ghana, often have no other choice than to seek treatment from traditional medical practitioners (TMP). The African ‘snakebite crisis’ is highlighted here using extensive complementary data from a community-based epidemiological study conducted by snake ecologists in the savanna zone of northern Ghana.

**Methodology and findings:** Our cross-sectional study included 1,000 residents and 24 TMPs in the Savelugu-Nanton District in northern Ghana between December 2008 and May 2009, and a 10-year (1999-2008) retrospective snakebite data from the district hospital. Variables tested included demography, human activity patterns, seasonality, snake ecology and clinical reports. Complementary data showed higher snakebite prevalence during the rainy season, and a hump-shaped correlation between rainfall intensity and snakebite incidences. Almost 6% of respondents had experienced a personal snakebite, whereas ∼60% of respondents had witnessed a total of 799 snakebite cases. Out of a total of 857 reported snakebite cases, 24 (∼2.8%) died. Highest snakebite prevalence was recorded for males in the age group 15-44 years during farming activities, with most bites occurring in the leg/foot region. Highest snakebite rate was within farmlands, most frequently caused by the Carpet viper (*Echis ocellatus*).

**Conclusion:** The relatively high community-based prevalence rate of ∼6%, and fatality rate of ∼3%, indicate that snakebites represent an important public health risk in northern Ghana. Based on the high number of respondents and long recording period, we believe these data truly reflect the general situation in rural Ghana and West Africa at large. We recommend increased efforts from both local and international health authorities to address the current snakebite health crisis generally compromising livelihoods and productivity of rural farming communities in West Africa.

**AUTHOR SUMMARY:** Snakebite envenomations cause tens of thousands of deaths and hundred thousands of injuries in many developing tropical countries each year, and sub-Saharan Africa represents an epitome of this ‘neglected tropical disease’. We present data spanning 10 years (1999-2008) which was collected over a six-month period, by applying different methodologies across a typical rural savanna community of northern Ghana. Our data corroborate previous findings from the region that snakebites constitute a serious public health threat, and that young and active farmers are particularly at risk, hence compromising both livelihoods and economic wealth of the people. We highlight that many interrelated factors involving both snake ecology and human behaviour in particular, are responsible for the high snakebite prevalence recorded. We conclude that our findings support increased and concerted efforts by both local authorities and state institutions to address the ongoing African snakebite crisis. Such interventions require the generation of more general baseline data on snake ecology and human behaviour, combined with education and information through public awareness campaigns. To achieve this, we recommend community-based stakeholder meetings involving the local people, traditional authorities, and public institutions working to address the persistent snakebite menace in this part of the world.

## INTRODUCTION

Snakebite envenomations constitute one of the most important human-wildlife conflicts, causing considerable yet largely insufficiently known magnitudes of socio-economical losses, morbidity and death [1]. Globally, out of >3,500 snake species, ∼600 are venomous, and ∼280 are considered medically important, causing a conservatively estimated >1.2 million snakebite envenomations causing ∼100.000 deaths and >400,000 cases of morbidity annually [1, 2, 3, 4]. Prevailing conservative estimates of the global burden of snakebite envenomations and fatalities are probably highly underrated as majority are based on conventional health facility reports, largely neglecting cases treated by local traditional medical practitioners (TMPs) [1, 5, 6]. Perhaps more realistic annual estimates indicate between 4.5-5.4 million snakebites, 1.8-2.7 million envenomations and up to 138,000 deaths [6, 7]. Snakebite envenomation is largely a disease of poverty, with developing countries in the tropics recording the highest rates of incidence, morbidity and mortality [4, 8, 9]. People engaged in farming, hunting, fishing and other rural activities are at highest risk, mostly bitten on their limbs during work [9, 10]. In many parts of sub-Saharan Africa, the high mortality and morbidity rates are attributed to increased vulnerability caused by both high work risk and exposure to diverse snake habitats, as well as poor infrastructure and limited access to appropriate medical treatment and health facilities [1, 4, 6]. Currently, an estimated ∼100 million people, particularly in southeast Asia and Africa, live in vulnerable areas with very high exposure to snake envenomation and lack of effective antivenom therapy [4]. The current ‘global snakebite crisis’ as a disease of poverty, has been termed misunderstood, underrated, ignored or neglected as a public health issue [5, 8, 11, 12], and has lately gained prominence as one of the most important ‘neglected tropical diseases’ [7].

In order to mitigate the inadequate health care and treatment of victims of snakebite envenomations, concerted international effort is essential to gather steady and inclusive data on the epidemiological nature of snakebites with socio-demographic and geographic dimensions as well as aspects of snake biology and ecology [4, 5, 12, 13]. Mapping of comprehensive datasets is imperative for understanding the dynamics of human-wildlife conflict such as snakebite vulnerability, and constitute the baseline information needed to provide adequate health facilities and supply of antivenom and other therapeutical innovations [1, 2, 6, 14]. However, inclusive community-based information is currently limited from many high-risk areas particularly in sub-Saharan Africa [15, 16]. There is therefore the need for detailed information from studies combining both field and hospital data [4, 6, 9]. Here, we present a comprehensive and complementary epidemiological dataset of snakebite envenomations from northern Ghana, comprising both household and TMP surveys as well as retrospective hospital reports covering the period 1999-2009. Apart from contributing baseline epidemiological snakebite data, our survey, conducted by snake ecologists, also targeted human-wildlife conflict dynamics, with the purpose of providing insight into measures for improved snakebite prevention and facilitation of effective therapeutic methods. We note that the vast majority of snakebite studies are undertaken solely by medical officers, and as such mainly focused on clinical-pharmaceutical, socio-demographic and epidemiological aspects, with much less attention to biology and ecology of humans as well as snakes. An additional objective of this paper was to apply an integrative approach involving snake ecology and common snakebite epidemiology, in order to increase our understanding of the complex human-wildlife conflict that snakebites truly represent. We consider that such holistic research foci are vital and urgently required by international funding agencies and national public health institutions in their joint efforts to address the ongoing global snakebite crisis [2, 4, 7, 10, 12, 13].

## METHODS

### Study area

The Savelugu-Nanton District, covering ∼2023 km^2^, is located in the Northern Region of Ghana. It is bordered by five other districts (Fig. 1), notably West Mamprusi (north), Karaga (east), Tolon-Kumbungu, and Tamale Metropolitan Area (west) and Yendi (south). Based on a 2010-population census count of 139,283 (female/male %-ratio = 51.5/48.5) and an annual growth rate of about 2.5% for northern Ghana [17], we estimated the total population of the district as ∼130,000 in 2008, translating to a mean density of ∼64 persons km^−2^. The district falls within the Guinea Savanna vegetation zone of northern Ghana, with a single-peak, erratic rainfall pattern, increasing rapidly from April to peak in August-September, then sharply decreasing during October, and ranging between ∼600 to ∼1,000 mm annually on average [18]. Mean daily temperatures are usually high, averaging 34 °C, with maximum of >42 °C and minimum of <15 °C [18]. Retrospective monthly rainfall data during 1999-2008 was obtained from the Ghana Meteorological Authority in Tamale. The sparsely-populated northern parts of the district have denser vegetation, mostly with regenerating woodlands, compared with the more urbanized south around the Tamale Metropolis characterised by more intensive farming, bush burning and tree felling for charcoal production. Many woodland tree species are drought-resistant and foliage is largely retained during the prolonged dry season (November-March).

**Figure 1.**
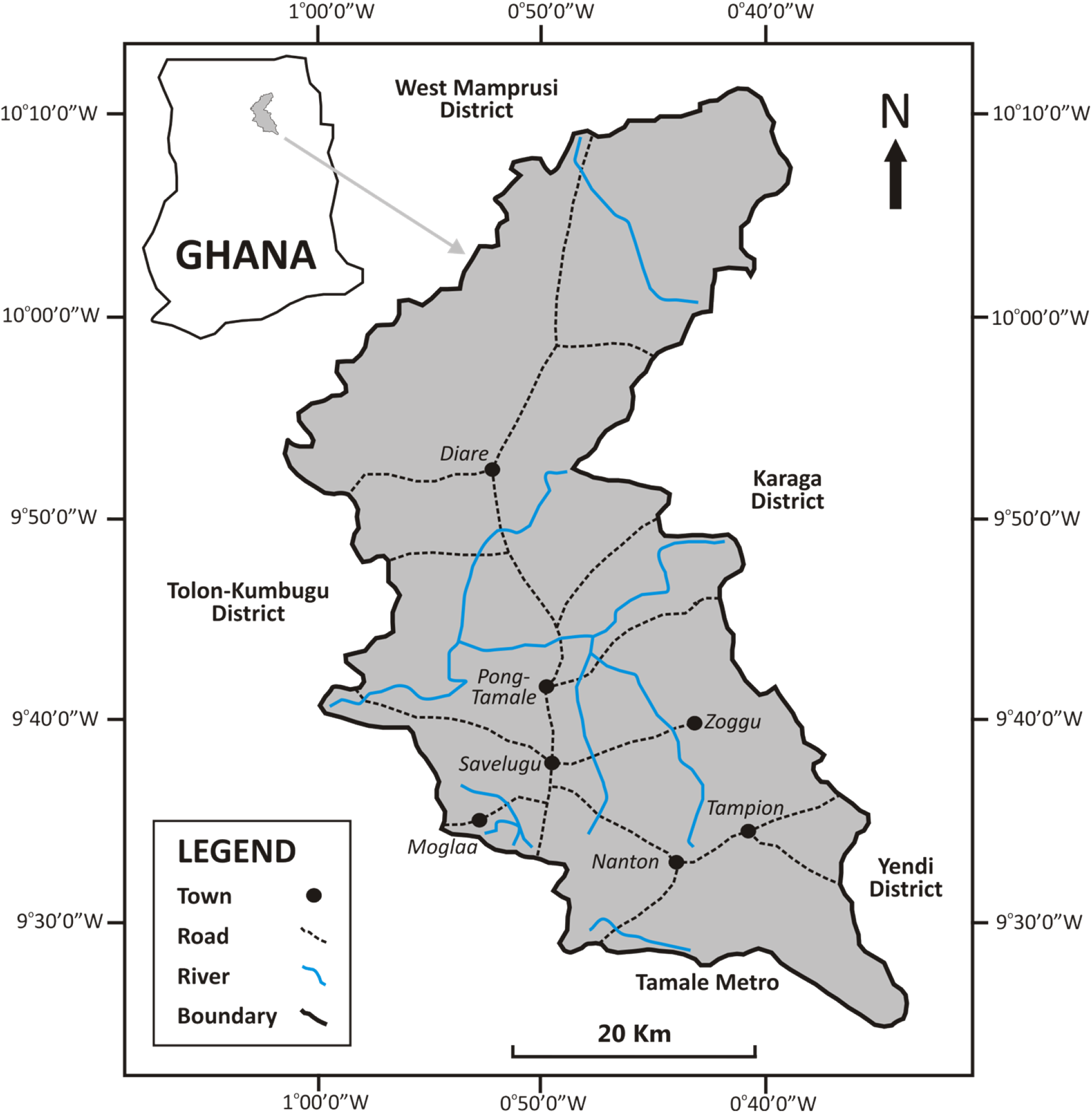
Map of the Savelugu-Nanton District in northern Ghana, showing major roads, rivers, and the seven selected townships for the study. Line drawings produced with CorelDraw™ version 12.

### Data collection

We conducted a cross-sectional respondent study between December 2008 and May 2009 in the seven most populated communities (= study sites) in the district (Fig. 1), including 1,000 community residents and 24 TMPs operating in these communities. Both residents and TMPs were subjected to detailed, independently administered, semi-structured questionnaires. Prior informed consent (PIC) was obtained from traditional leaders and authorities in each community after explanation of the scope, purpose and procedure of the study [19]. Such independent administration of interview protocols and PIC enhance the reliability and inclusiveness of respondent information and hence both the quantity and quality of informant data [20, 21]. The selection of residents was stratified randomly across three age groups, each with gender equality; >30 years (400); 15-30 years (300); <15 years (300). The number of residents sampled from each of the seven study sites reflected community size and were; Savelugu (200), Diarre and Pong-Tamale (150 each), Nanton, Zoggu, Moglaa and Tampion (125 each). Within each of the seven communities, respondents were randomly selected among households, inclusively conforming to the gender-age stratification criteria. The 24 TMPs were selected opportunistically using ‘Snowball sampling’ [19]. Complementing the community-based data collection was a retrospective study of reported cases of snakebites over a 10-year period (1999-2008) that were obtained from the Savelugu District Hospital, the largest district hospital managing snakebite victims. Patient data was analysed anonymously, after obtaining formal approval from the Savelugu District Hospital authorities.

### Data analysis

Informant data obtained from questionnaires were analysed using Microsoft Excel™ and GraphPad™ version 5.01. Frequencies of occurrence (%) in each respondent category were compared among different sub-groups, and associations tested for statistical differences with Fisher’s exact test for 2×2 contingency tables (two sub-groups compared) or χ^2^ *G*-test for three or more sub-groups compared (2×3 contingency tables). Associations between rainfall and snakebite incidences were determined using both GLM and polynomial regression with Pearson’s (*r*) or Spearman rank (*r_s_*) correlation coefficients. Significance level was determined at p < 0.05.

## RESULTS

### Household survey

#### Educational level of respondents

Majority (518) of 1,000 respondents (∼52%) had no formal education whereas ∼44% had only basic primary education (7-9 years of schooling). Only ∼4% had additional secondary education (10-12 years), with <0.5% having tertiary education (>12 years) only (Table 1). Hence, respondents were primarily made up of either illiterates or people with modest schooling.

**Table 1.**
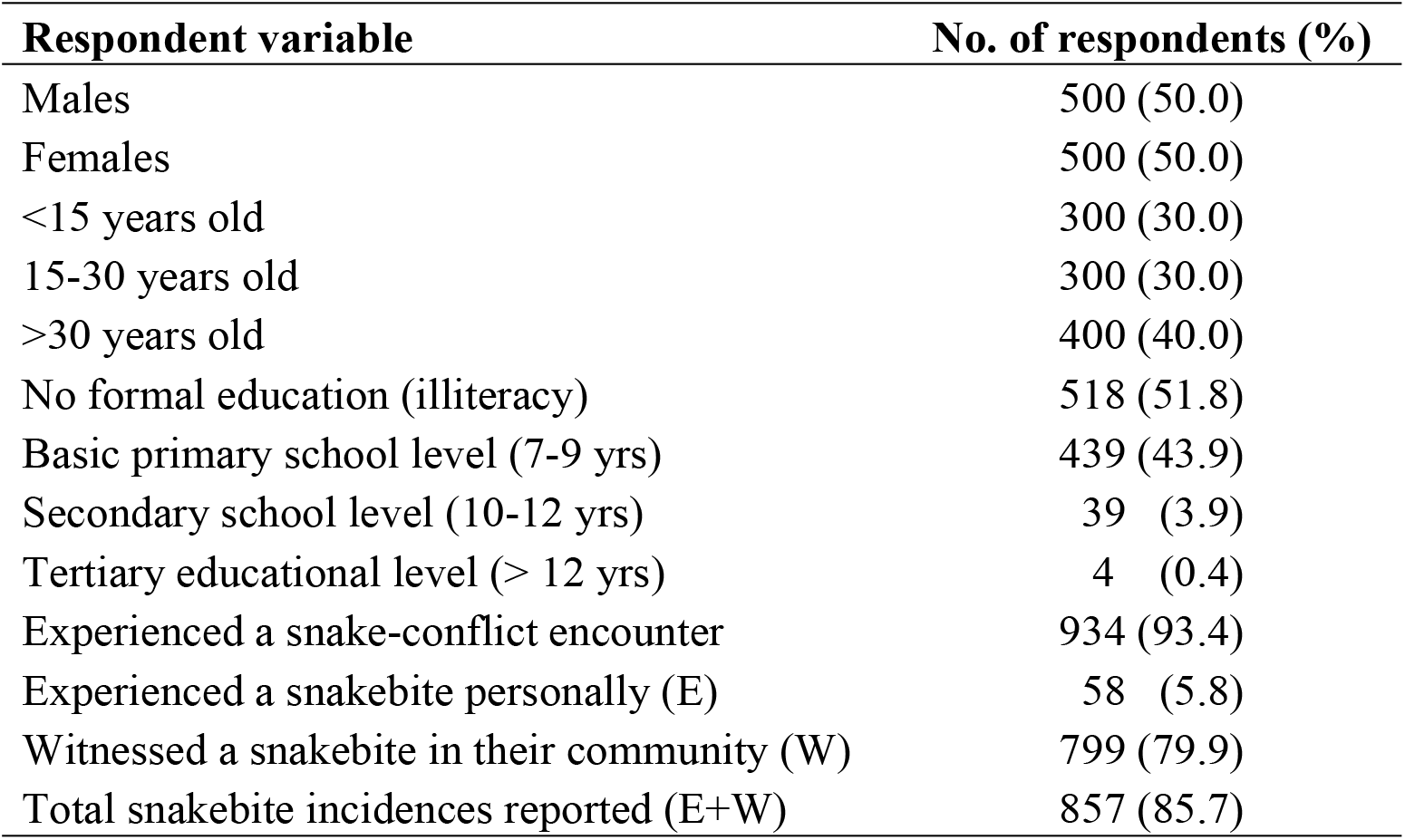
Basic socio-demographic characteristics and snake-conflict encounter statistics of the respondent community population (n = 1,000 residents) comprising seven communities surveyed in the Savelugu-Nanton District of northern Ghana (Dec 2008-May 2009).

| Respondent variable | No. of respondents (%) |
| --- | --- |
| Males | 500 (50.0) |
| Females | 500 (50.0) |
| <15 years old | 300 (30.0) |
| 15-30 years old | 300 (30.0) |
| >30 years old | 400 (40.0) |
| No formal education (illiteracy) | 518 (51.8) |
| Basic primary school level (7-9 yrs) | 439 (43.9) |
| Secondary school level (10-12 yrs) | 39 (3.9) |
| Tertiary educational level (> 12 yrs) | 4 (0.4) |
| Experienced a snake-conflict encounter | 934 (93.4) |
| Experienced a snakebite personally (E) | 58 (5.8) |
| Witnessed a snakebite in their community (W) | 799 (79.9) |
| Total snakebite incidences reported (E+W) | 857 (85.7) |

#### Snakebite prevalence: comparing gender and age of victims

Considerably more males (∼62%) than females (∼38%) were reportedly bitten by snakes (n = 58 cases), although the difference was not statistically significant (Fisher’s exact test; p = 0.0778, df = 1), at the 5% level (Table 2). With regards to age, significantly (χ^2^ = 14.616; p = 0.00067, df = 2) more victims were in the age group >30 years (∼64%), whereas the group of 15-30 years (∼17%) and <15 years (∼19%) were similar in prevalence to snakebites (Table 2).

**Table 2.** Associations of reported personal snakebite incidences (n = 58) with gender and age-group of 1,000 community members (male/female = 500 each) interviewed in the Savelugu-Nanton District of northern Ghana (Dec 2008-May 2009).

| Variable | Sub-group variable | Frequency (%) | Test-statistics | p-value |
| --- | --- | --- | --- | --- |
| Gender | Male | 36 (62.1) | Fisher's exact test (2×2) | 0.0778 |
|  | Female | 22 (37.9) |  |  |
| Age | <15 years | 11 (19.0) | $\chi^2$ G-test (2×3) = 14.616 | 0.00067 |
|  | 15-30 years | 10 (17.2) |  |  |
|  | >30 years | 37 (63.8) |  |  |

#### Personal encounters, interactions and snakebites

As many as 934 (∼93%) respondents had encountered snakes during their lifetime and hence been exposed to considerable risk of snakebites, and 58 respondents (∼6%) claimed to have been bitten by snakes, irrespective of actual envenomation or hospitalisation (Table 1). Of the 934 snake encounters (Table 3) majority encountered snakes on their farms (604; ∼65%), but also in the bush (∼16%) or in their homes (∼11%). Considerably lower encounter frequencies were attributed to roads and footpaths (∼6%), school facilities (∼2%) and open urban drains (<1%). Snake encounters were predominant during afternoon (∼50%) and morning (∼40%) hours, particularly during the rainy season, accounting for ∼72% of yearly records by respondents (Table 3).

**Table 3.** Characteristics of personal snake-conflict encounters reported (n = 934 incidences) among 1,000 community members interviewed in the Savelugu-Nanton District of northern Ghana (Dec 2008-May 2009).

| Respondent variable | Sub-group variable | Encounters (%) |
| --- | --- | --- |
| Encountering habitat location | Farmlands | 604 (64.7) |
|  | Bushes | 145 (15.5) |
|  | House and yards | 100 (10.7) |
|  | Paths and roads | 60 (6.4) |
|  | School facilities | 19 (2.0) |
|  | Open city drains | 6 (0.6) |
| Encountering time of the day | Afternoon | 471 (50.5) |
|  | Morning | 372 (39.8) |
|  | Evening and night | 91 (9.7) |
| Encountering season of the year | Rainy season | 669 (71.6) |
|  | Dry season | 265 (28.4) |

#### Knowledge, awareness and perception of other snakebite victims

A total of 604 (∼60%) respondents provided knowledge about 799 snakebite cases from their respective communities in total (Table 4). Thus, the total recorded cases of snakebites, including personal experiences (n = 58) was 857, with a total of 24 (∼2.8%) reported deaths (Table 1, 5).

**Table 4.**
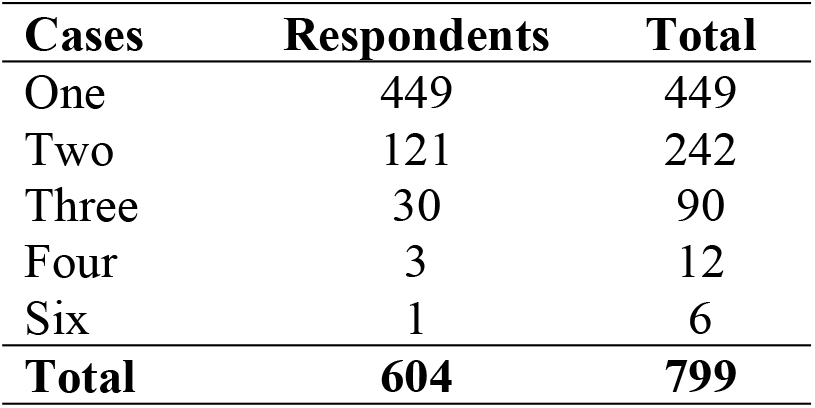
Snakebite cases reported by each of 604 respondents out of 1,000 community members interviewed in the Savelugu-Nanton District of northern Ghana (Dec 2008-May 2009).

**Table 5.**
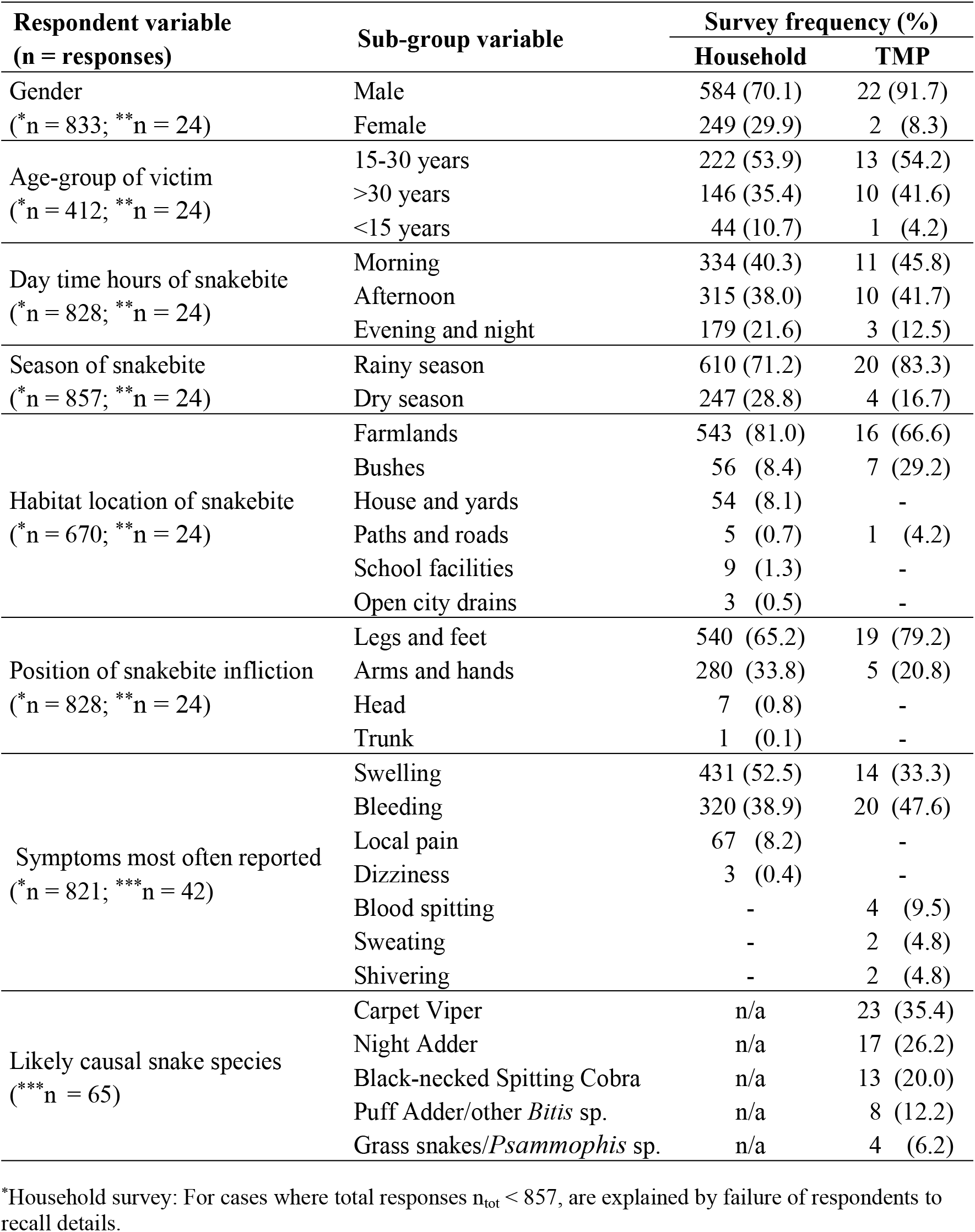
Comparison of perceptual characteristics of snakebite victims and incidences based on household surveys (*n_max_ = 857 cases) and traditional medical practitioners (TMP; **n = 24 respondents; ***n = 42 and 65 cases), respectively among 1,000 community members and 24 TMPs, interviewed in the Savelugu-Nanton District of northern Ghana (Dec 2008-May 2009).

| Respondent variable<br>(n = responses) | Sub-group variable | Survey frequency (%) |  |
| --- | --- | --- | --- |
|  |  | Household | TMP |
| Gender<br>(*n = 833; **n = 24) | Male | 584 (70.1) | 22 (91.7) |
|  | Female | 249 (29.9) | 2 (8.3) |
| Age-group of victim<br>(*n = 412; **n = 24) | 15-30 years | 222 (53.9) | 13 (54.2) |
|  | >30 years | 146 (35.4) | 10 (41.6) |
|  | <15 years | 44 (10.7) | 1 (4.2) |
| Day time hours of snakebite<br>(*n = 828; **n = 24) | Morning | 334 (40.3) | 11 (45.8) |
|  | Afternoon | 315 (38.0) | 10 (41.7) |
|  | Evening and night | 179 (21.6) | 3 (12.5) |
| Season of snakebite<br>(*n = 857; **n = 24) | Rainy season | 610 (71.2) | 20 (83.3) |
|  | Dry season | 247 (28.8) | 4 (16.7) |
| Habitat location of snakebite<br>(*n = 670; **n = 24) | Farmlands | 543 (81.0) | 16 (66.6) |
|  | Bushes | 56 (8.4) | 7 (29.2) |
|  | House and yards | 54 (8.1) | - |
|  | Paths and roads | 5 (0.7) | 1 (4.2) |
|  | School facilities | 9 (1.3) | - |
|  | Open city drains | 3 (0.5) | - |
| Position of snakebite infliction<br>(*n = 828; **n = 24) | Legs and feet | 540 (65.2) | 19 (79.2) |
|  | Arms and hands | 280 (33.8) | 5 (20.8) |
|  | Head | 7 (0.8) | - |
|  | Trunk | 1 (0.1) | - |
| Symptoms most often reported<br>(*n = 821; ***n = 42) | Swelling | 431 (52.5) | 14 (33.3) |
|  | Bleeding | 320 (38.9) | 20 (47.6) |
|  | Local pain | 67 (8.2) | - |
|  | Dizziness | 3 (0.4) | - |
|  | Blood spitting | - | 4 (9.5) |
|  | Sweating | - | 2 (4.8) |
|  | Shivering | - | 2 (4.8) |
| Likely causal snake species<br>(***n = 65) | Carpet Viper | n/a | 23 (35.4) |
|  | Night Adder | n/a | 17 (26.2) |
|  | Black-necked Spitting Cobra | n/a | 13 (20.0) |
|  | Puff Adder/other *Bitis* sp. | n/a | 8 (12.2) |
|  | Grass snakes/*Psammophis* sp. | n/a | 4 (6.2) |

\*Household survey: For cases where total responses n<sub>tot</sub> < 857, are explained by failure of respondents to recall details.

Of the 857 snakebite cases, majority (∼70%) of victims were males, predominantly in the 15-30 years (∼54%) and >30 years (∼35%) age groups. Farmlands were the most frequently reported locations (∼81.0%) for snakebite, primarily during the rainy season (∼71%), in the mornings (∼40%) and afternoons (∼38%) (Table 5). Snakebites occurred mainly on the extremities, with lower limbs (legs and feet; ∼65%) and upper limbs (arms and hands; ∼34%) being the most vulnerable. Swelling (∼53%) and bleeding (∼39 %) were the commonest symptoms reported (Table 5).

### Traditional medical practitioner survey

#### Knowledge and perception of snakebite victims and snake culprits

The 24 TMPs reported that majority of patients were males (∼92%), in the age groups >30 years (∼54%) and 15-30 years (∼42%), most often bitten on farmlands (∼67%) and bushes (∼29%), during the rainy season (∼83%), in the mornings (∼46%) and afternoons (∼42%) hours (Table 5). According to the TMPs, most vulnerable are legs and feet (∼79%), followed by the arms and hands (∼21%), with bleeding (∼48%) and swelling (∼33%) as most frequently reported symptoms (Table 5). The general snake descriptions provided by the TMPs indicated that three snake species were most likely involved in ∼81% of all cases reported; Carpet Viper *Echis ocellatus* (∼35%), Night Adder *Causus maculatus* (∼26%), and Black-necked Spitting Cobra *Naja nigricollis* (∼20%).

### Hospital survey

#### Snakebite prevalence: comparing gender and age of victims

Based on the 10-year (1999-2008) retrospective records of snakebite cases (n = 450) at the main district hospital, significantly more males, particularly in the 15-44 years age group (χ^2^ G-test = 29.56, p < 0.00001, df = 2), were bitten than females, as well as males in the <15 years and >44 years age groups (Table 6).

**Table 6.**
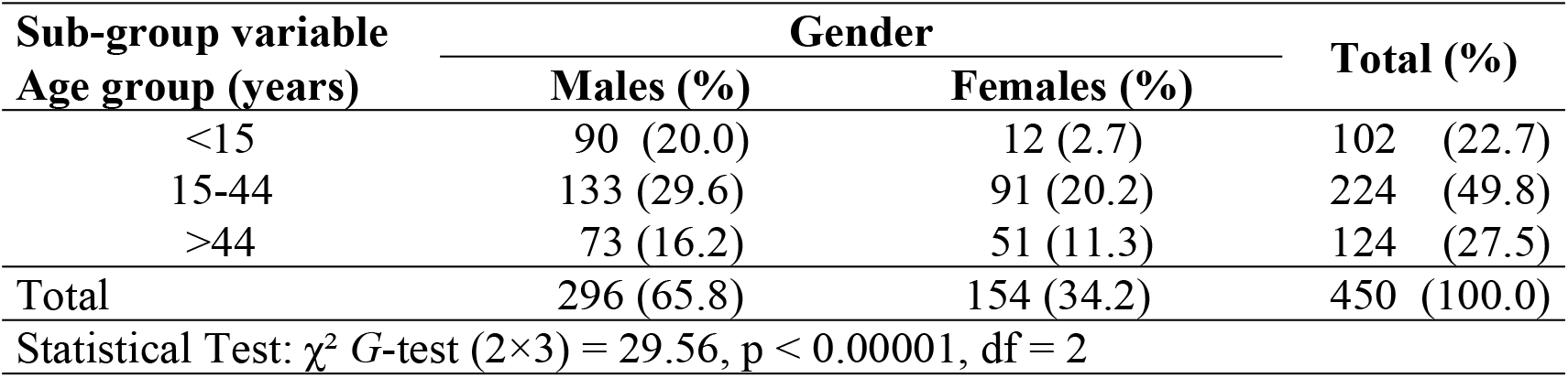
Statistically-tested associations of snakebite cases (n = 450) with gender and age-group reported at the Savelugu-Nanton District Hospital in northern Ghana (1999-2008).

Across all three age groups, males dominated in number of reported cases, whereas the females constituted relatively fewer cases in the youngest age group (2.7%) but 4-8 times more in the two older groups (Table 6). For both males and females, the 15-44 years group dominated in almost 50% of all cases reported. These results are largely consistent with age-gender trends of personal snakebite cases (n = 58) reported among 1,000 respondents in the 2008-2009 household survey, with dominance of males across all age groups, and in particular for the >30 years age group (Table 2). In summary, the most and least vulnerable to snakebites were young to middle aged men and younger girls respectively.

#### Correlations between rainfall and snakebites for the period 1999-2008

By comparing concurrent data sets of mean monthly rainfall with monthly records of snakebites reported to the district hospital in 1999-2008 (Fig. 2), we were able to evaluate the perception reported among community members that snakebites were more prevalent during the rainy season (Table 3, 5). The 10-year plot of rainfall-snakebite monthly means depicts a clear unimodal rainfall curve with a single peak from July to September, whereas snakebite incidences display a trimodal pattern with peaks in March, June-July and October (Fig. 2). Hence, the largely overlapping rainfall-snakebite trend demonstrates a positive association between rainfall and snakebite frequency, in line with respondent (both victims and TMPs) perceptions (Table 3, 5). We tested this apparent positive association by plotting 10-years pair-wise data of mean monthly rainfall with mean monthly snakebite cases, and found a weak statistically significant (*r_s_* = 0.587, p = 0.0446, two-tailed, n = 12) positive linear correlation between the two variables (Fig. 3). However, the correlation for GLM-regression using Pearson’s correlation coefficient was not significant (*r* = 0.521, p = 0.0822, two-tailed, n = 12). We also performed a 2^nd^ degree polynomial regression which showed a higher level of significance (*r* = 0.652, p = 0.022, two-tailed, n = 12) compared with the GLM-regression, thereby indicating a hump-shaped association pattern (Fig. 3). Thus, although snakebite rates are generally linked to rainfall levels, extreme amounts of rain appear to reduce this positive correlation.

**Figure 2.**
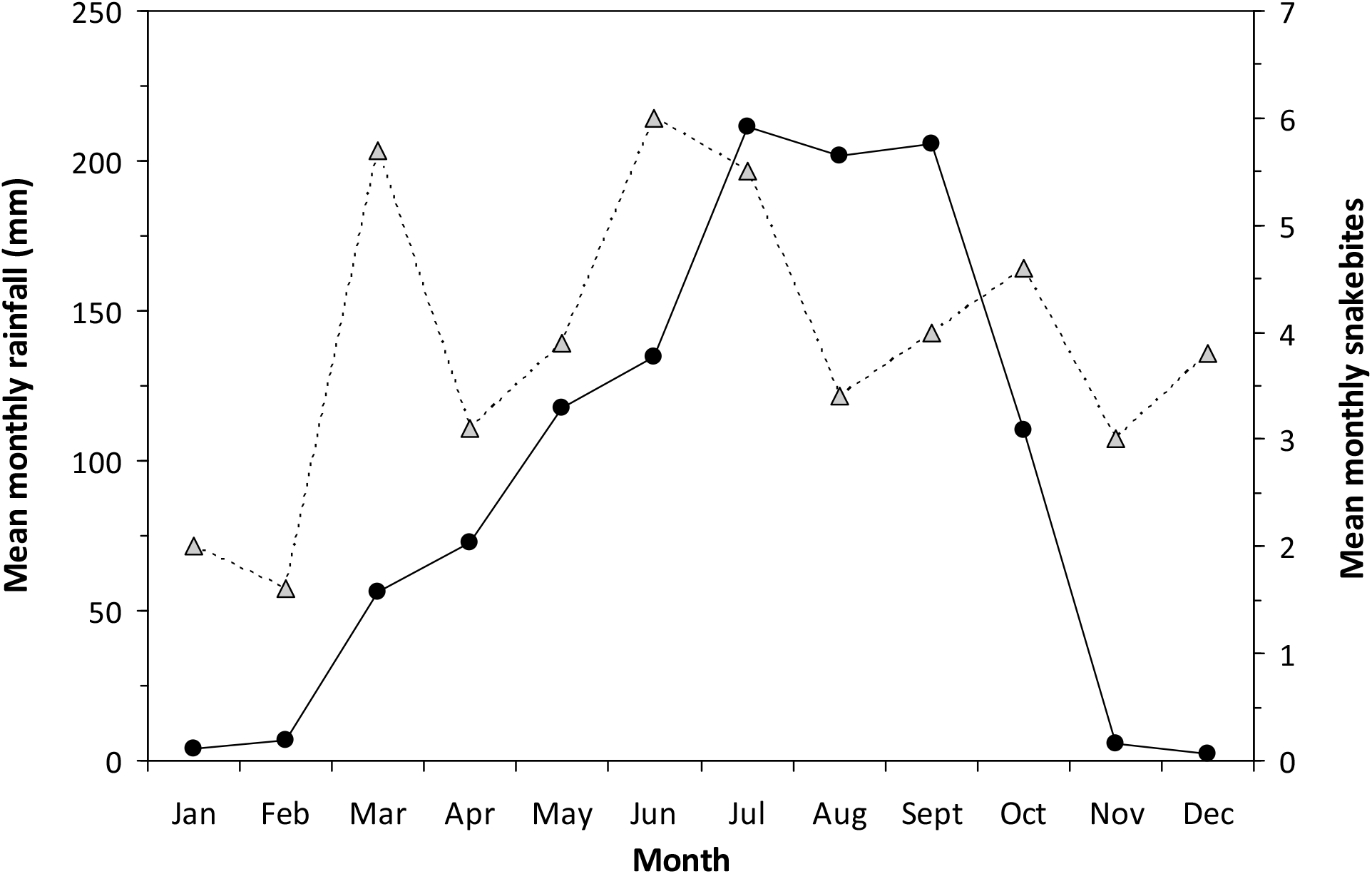
Mean monthly precipitation (black circles and solid line) and mean monthly number of snakebite incidences (grey triangles and broken line) recorded at the Savelugu District Hospital (1999-2008).

**Figure 3.**
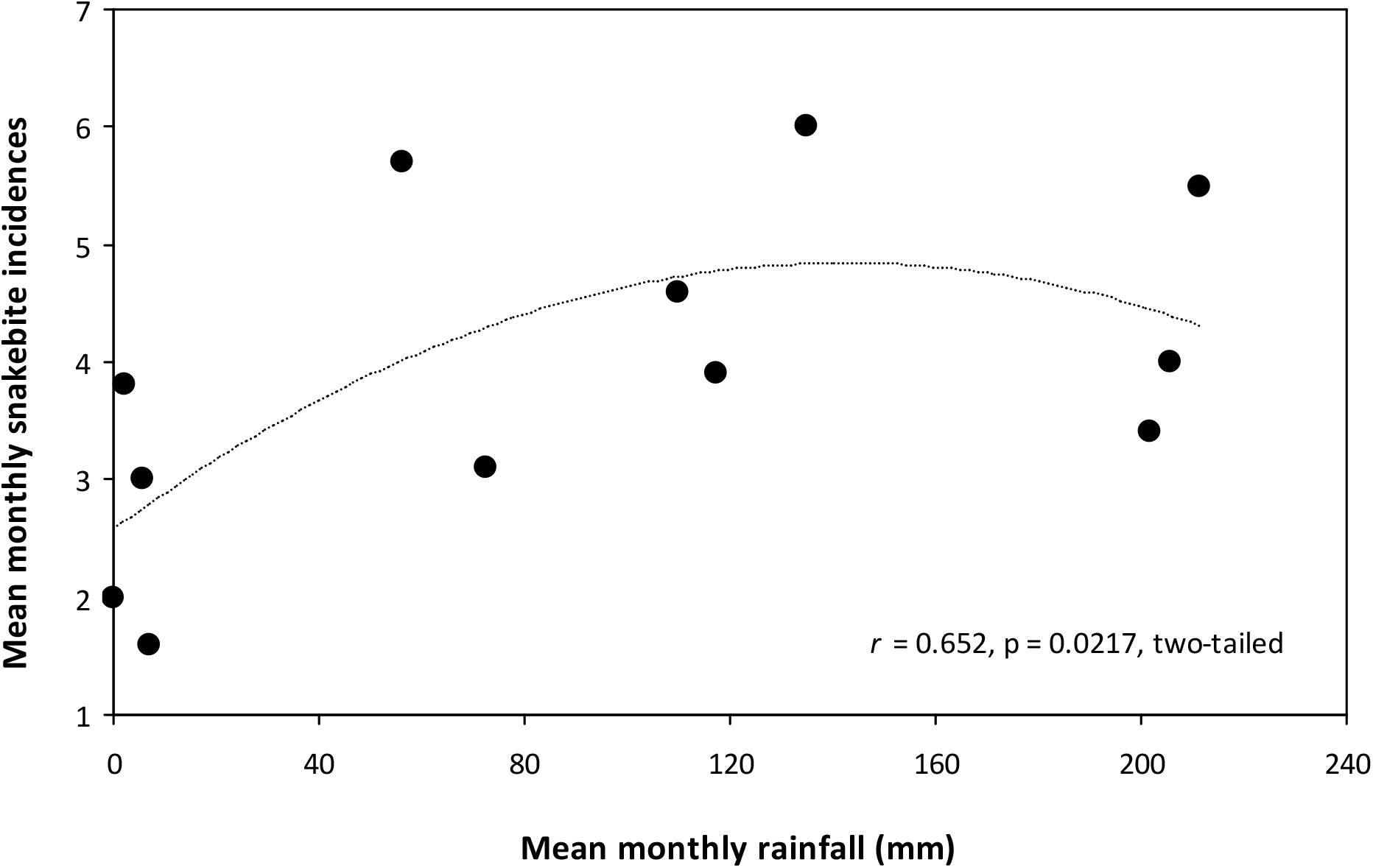
Correlation between mean monthly precipitation and mean monthly number of snakebite incidences recorded at the Savelugu District Hospital (1999-2008).

## DISCUSSION

Our study applied an extensive set of complementary epidemiological data from different community sectors, by targeting various demographic sub-groups as well as both traditional and non-traditional management of snake envenomations at the district level. With a large sample size of 1,024 respondents covering both dry and wet seasons, and a retrospective 10-year bivariate data sample of snakebite incidences and rainfall patterns, our study represents an important contribution to epidemiological snakebite studies for this region. Previous epidemiological studies have largely been limited to short-term data series with lower sample sizes of predominantly retrospective hospital records for Ghana [22, 23, 24] and sub-Saharan Africa [25, 26, 27, 28, 29], where complementary or nationwide long-term published studies remain scarce [1, 16, 30, 31]. Estimates of regional or national snakebite burdens based exclusively on hospital records inevitably neglect many snakebite cases treated at home or by TMPs [1, 4, 16, 30]. In contrast, our study provided a rare opportunity to test consistencies in epidemiological baseline information gathered from different sections of a typical rural savanna community in West Africa [16], regarding general demography and human-snake conflict, as well as snake toxicology and ecology. In order to enhance advances in public health capacity to address snakebite envenomation, it is important to provide inclusive and holistic evaluations of implicating factors across the whole community and over long periods of time, including ecological aspects [2, 4, 7, 30]. Specifically, it is important to ask where, when and why snakebites occur, and to identify and evaluate pertinent factors that preclude or facilitate their occurrence, treatment and prevention [1, 4, 10, 12].

Our study confirms the general assertion that snake envenomations leading to significant levels of morbidity and mortality are symptomatic of rural deprivation and poverty in sub-Saharan Africa, where educational levels are pronouncedly low within farming communities subject to limited infrastructure and mobility [4, 8, 9, 12]. Our respondents were primarily made up of either illiterates or people with modest schooling, a demographic feature characteristic of the mostly deprived rural areas of northern Ghana, and a tendency particularly pronounced among women and the elderly. We located 24 TMPs in seven randomly-chosen communities, demonstrating that the TMP-inhabitant ratio of at least 1:5,000 (∼24/130,000) is ∼10 times lower than the regional doctor-patient ratio for northern Ghana in 2009 [32]. As the Savelugu-Nanton District road network was fairly well developed during our study, we attribute the high patronage of TMPs mainly to low educational level and income, limited access to orthodox medical facilities as well as high costs or lack of sufficient and effective antivenom administration. This TMP-seeking behaviour is widespread in Asia [33] and Africa [16, 34, 35, 36] where herbal extracts of diverse medicinal plant assemblages [37] are often either culturally preferred or are the only affordable or alternative therapeutic treatment available among low-income rural communities with high levels of illiteracy and strong belief in the supernatural [16, 19, 38, 39, 40, 41].

Based on the 1,000-respondent household survey, we conservatively estimated a snakebite prevalence of ∼6%, and a mortality rate of ∼3%, translating to respectively ∼7,500 snakebite incidences and ∼200 deaths at the district level, for a total district population estimated at ∼130,000 in 2008. We recognise that our incidence and mortality data are based on unknown recollection periods of each informant, and we can therefore not explicitly translate these district level figures into annual snakebite and mortality rates. However, assuming a recollection period of ∼10 years retrospectively for an average respondent, the estimated annual snakebite cases and fatalities at district level may probably reach ∼750 and ∼20 respectively. A longer average respondent recollection period is unlikely as both young and old respondents exhibited limited memory beyond 10 years. Moreover, if such a period was ∼5 years, it would translate to a double of cases and fatalities, which likewise appears unlikely. The mean monthly number of cases reported to the largest district hospital was 3.9 over the 10-year period (1999-2008), which equals an average of 46.6 cases yearly (Fig. 2). Based on hospital records, the whole district therefore probably has a burden of at least 50 reported cases yearly, and probably as much as double if the reporting percentage is up to 50% as estimated in similar studies [12, 16, 41]. Our upper-lower limits of annual snakebite envenomations during the 10-year period therefore range from 100-750, equivalent to 75-575 per 100,000 inhabitants, with a mortality rate of 2-17 per 100,000. The population of Savelugu-Nanton District was ∼130,000, constituting ∼6% of the population of the Northern Region of Ghana in 2008 [17], so we conservatively estimate about 1,700-12,500 envenomations with about 50-350 fatalities annually (mortality rate of ∼3%) in this part of the country. This figure makes up about 20-95% of a Ghana national estimate of about 250-375 [40], and is within the range of current available estimates for the sub-region [1, 9] and other estimates from northern [23] and central [22, 42] Ghana, but 5-10 times higher than southwest Ghana [24].

The human deprivation, psychosocial despair, and loss of income resulting from snakebite morbidity and mortality [7, 35, 40], are clearly demonstrated by our complementary data from the northern part of the savanna zone in Ghana, where annual snakebite burdens are the most severe across the entire country. Our data also indicate that the vast majority of snakebites are never reported to conventional medical treatment facilities but managed by TMPs, home treatment or are untreated, sometimes with fatal consequences [16, 24, 43]. Long delays in antivenom treatment often lead to severe cases of morbidity (e.g. amputations) or death, even if such treatment is administered correctly [22, 42, 44, 45]. It is therefore important to ascertain the extent of both untreated and delayed treatments in order to address these clinical and epidemiological shortcomings in the management of snakebites in remote rural parts of Ghana and sub-Saharan Africa [2, 13, 15, 40, 42]. We hope our snakebite data analyses from the Savelugu-Nanton District, a typical northern savanna community in Ghana, will aid in this direction.

We consistently found that young to middle-aged men (age group = 15-44 years) were most at risk of snakebites, corroborating similar findings from Ghana [22, 23, 24], other parts of West Africa [15, 25, 26, 27, 44, 46, 47] and Africa in general [30, 34, 48, 49, 50]. Adolescent and young men in their twenties are among the most active and adventurous, albeit least cautious, section of rural African people [24, 27, 44, 51]. They expose themselves to snake encounters by risky behaviour during land clearing, harvesting, bush and charcoal burning, hunting and commuting on foot during dark or early morning hours with impaired visibility in dim light and dense vegetation [24, 47, 50]. Minors and the elderly who are traditionally home-bound, are less involved in farming activity, and are generally more cautious in their behaviour, which possibly explains their much lower snakebite vulnerability [31]. Some girls and the elderly were however bitten at home or in school facilities, indicating that everyone is at risk of snakebites [7, 24, 31, 43, 51, 52]. Contrary to our findings, some studies in Africa actually report relatively higher snakebite prevalence in children, including girls, probably as a result of higher child-labour engagement or poorer parental care and supervision in such areas [23, 31, 50, 51, 53, 54, 55, 56].

Our data from all three respondent groups indicated that farmlands and bushes, and to a lesser extent, residential areas and roads are typical habitats for snakebite, as corroborated by several other studies in rural Africa [16, 31, 38, 50, 51]. Likewise, majority of bites occur during the peak farming periods (e.g. shea nut, millet and yam) when land is prepared for planting (March-April), maintained by weeding (May-June), and harvested (June-October) [23, 47, 51]. Our long-term bivariate rainfall-snakebite data indirectly reflect the strong link between peak seasons of farming and rainfall, partly because artificial water sources are largely unavailable to the small-scale peasant farmer in northern Ghana and other parts of arid savanna zones in Africa. However, even though rainfall and snakebites are generally positively-correlated, our data also suggest that very intense rainfall reverses the trend, possibly caused by decreased activity of both snakes and humans as well as lower snakebite reporting rates due to the reduced mobility from flooding and erosion of basic road infrastructure. Although the dry season recorded lower snakebite rates, farmers are also at risk during this period, often being bitten during bush or charcoal burning, hunting, harvesting firewood, and at home during reduced farming activity in November-April [22, 47, 50].

As agricultural mechanisation is particularly low in northern Ghana, predominantly manual activities using the limbs expose farmers to snakes concealed in vegetation, soil and crops. This may explain why those extremities are the most prone to snakebites [23, 44, 47, 51]. Farming activities primarily take place during morning or late afternoon in order to avoid excessive sun exposure and heat during midday. Such factors also result in the poikilothermic snakes seeking warm areas during morning hours and shade in the hot midday and early afternoons. Nocturnal snakebites were mostly confined to homes and roads, a pattern found in many other rural areas of Africa, and attributed to walking barefoot, or inappropriate footwear and poor domestic lighting conditions [25, 31, 45, 51].

Snakebite prevalence, risk and rates are determined by several interrelated factors pertaining to both snakes and humans, notably the density, activity and behaviour of the snake culprits and their victims. Snake density appears the most important factor, as the commonest species are also often the most prevalent as culprits [23, 46]. Snake density is related to habitat type and human disturbance, as well as food sources and hiding places. It is also inversely related to human density in many parts of sub-Saharan Africa [27, 30, 46], and therefore directly related to both destruction of suitable habitats as well as human persecution, especially in most African cultures with great dislike of snakes. Snakebite prevalence is lower in urban centres devoid of natural vegetation, and where snakes are at higher risk of being detected and killed [30]. Our data were in line with this inverse human-snake density-relationship, as evidenced by lower bite rates closer to Tamale, the district capital with >300,000 inhabitants.

Apart from densities of both humans and snakes, activity patterns and behaviour are two other important factors highly correlated with habitats, daytime hours and season. During rains, in which farming activities peak in the savanna zone, snakes often synchronise their breeding periods with high prey abundances [57, 58]. Likewise, density and activity of prey such as amphibians, birds, lizards, murid rodents and other snakes increase during rains, due to prey availability (mainly invertebrates) increasing along with the flourishing vegetation. The number of active farmers, combined with actively-hunting and breeding snakes tends to increase human-snake encounters, often resulting in bites and envenomations [46]. Similarly, the density-activity influence on human-snake encounters and the behaviour of the snake culprits as well as victims are important factors. Risky behaviour during farming, hunting and firewood collection (e.g. using unprotected limbs to grasp, cut, dig, lift and pick tools, foods and other important farm-related items), increases the probability of unintendedly provoking and even attacking snakes [31, 51]. In response, the disturbed snakes may vigorously defend themselves either by hissing, inflating hoods, whipping tails, venom spitting or biting. Certain species appear more docile and reluctant to strike even if trodden on (e.g. Gaboon Viper *Bitis gabonica* and Phillips Grass Snake *Psammophis phillipsi*) while others tend to be extremely aggressive and prone to strike at the least provocation or threat (e.g. Black-necked Spitting Cobra *Naja nigricollis* and Carpet Viper *Echis ocellatus*). Some species retreat long before humans are even close, whereas others stay put and motionless until it is too late to detect their presence before they strike. Highly mobile species may enter houses, garages and even cars in search of prey or warm places (e.g. cobras and small vipers), while other slow-moving species (e.g. large vipers) avoid such human-trafficked areas which pose great human-detection risks. As in the majority of West African studies, and as recorded from all sections of our study population, three snake species were by far the commonest snakebite culprits; Carpet Viper, Night Adder *Causus maculatus* and Black-necked Spitting Cobra [16, 23, 26, 35, 40, 43, 45, 46]. These three species were also the commonest found in our study area, as shown by extensive ecological censuses [59]. Polyvalent antivenom of these species in particular, is therefore essential in this part of northern Ghana [22, 23, 42].

## CONCLUSION

Our community-based complementary data clearly demonstrate the gravity of snakebite envenomations in the savanna zone of northern Ghana, and the relatively high burden of incidence, morbidity and mortality. These findings cannot be overemphasised with respect to their negative implications for public health, agricultural productivity, social welfare and economic growth. In order to address the shortcomings of adequately-trained personnel, effective treatment with antivenom, and other efficient therapy, we recommend concerted efforts from both conventional and traditional medical practitioners to monitor, map and analyse snakebite incidences at district levels, including scientific efficacy testing of complementary therapy to expensive and unavailable polyvalent antivenoms. Additionally, community members, particularly the youth, should be sensitised on risky snakebite behaviour, snake biology and measures to prevent and minimise the likelihood of snakebites. Such interventions could be achieved through community meetings, education and awareness sessions, and steady contact in the field with public health authorities liaising with traditional rulers and TMPs.

## ACKNOWLEDGEMENTS

We would first like to thank the local communities that participated in the survey, notably the elders, traditional rulers and medicinal practitioners, as well as staff at the Savelugu District Hospital and the Meteorological Survey Department in Accra. Special appreciation goes to Mohammed Mukhtar, Imoro Abdul Rashid, Issah Shaban, Abubakari Abdul Fatawu and Yakubu Abdul Latif.

## SUPPORTING INFORMATION

S1 Checklist. STROBE checklist.

Checklist of items for this observational cross-sectional study.

## REFERENCES

1. Kasturiratne A, Wickremasinghe AR, De Silva N, Gunawardena NK, Pathmeswaran A, Premaratna R, Savioli L, Lalloo, DG., de Silva, HJ. The global burden of snakebite: A literature analysis and modelling based on regional estimates of envenoming and deaths. PLoS Medicine. 2008; 5(11):1591–604.

2. Gutiérrez JM, Warrell DA, Williams DJ, Jensen S, Brown N, Calvete JJ, Harrison RA (2013) The need for full integration of snakebite envenoming within a global strategy to combat the neglected tropical diseases: The way forward. PLoS Neg. Trop. Dis. 7(6): e2162. doi:10.1371/journal.pntd.0002162

3. Figueroa A, McKelvy AD, Grismer LL, Bell CD, Lailvaux SP (2016) A species-level phylogeny of extant snakes with description of a new colubrid subfamily and genus. PLoS ONE 11(9): e0161070. doi:10.1371/journal.pone.0161070.

4. Longbottom J, Shearer FM, Devine M, Alcoba G, Chappuis F, Weiss DJ, Ray SE, Ray N, Warrell D, de Castañeda RR, Williams DJ, Hay SI, Pigott DM (2018) Vulnerability to snakebite envenoming: a global mapping of hotspots. The Lancet 392: 673–684. Published online at http://dx.doi.org/10.1016/S0140-6736(18)31224-8.

5. Gutiérrez JM, Theakston RDG, Warrell DA (2006) Confronting the neglected problem of snake bite envenoming: The need for a global partnership. PLoS Med 3(6): e150. DOI: 10.1371/journal.pmed.0030150.

6. Gutiérrez JM, Williams D, Fan HW, Warrell DA (2010) Snakebite envenoming from a global perspective: Towards an integrated approach. Toxicon 56: 1223–1235. doi:10.1016/j.toxicon.2009.11.020.

7. World Health Organisation (2017) Global snakebite burden. Report by the Director-General (EB142/17). Provisional agenda item 4.1. (18 Dec 2017).

8. Cruz LS, Vargas R, Lopes AA (2009) Snakebite envenomation and death in the developing world. Ethnicity & Disease 19: S1-42 – S1-46.

9. Harrison RA, Hargreaves A, Wagstaff SC, Faragher B, Lalloo DG (2009) Snake Envenoming: A disease of poverty. PLoS Negl Trop Dis 3(12): e569. doi:10.1371/journal.pntd.0000569.

10. Chippaux J-P, Akaffou MH, Allali BK, Dosso M, Massougbodji A, Barraviera B. The 6th international conference on envenomation by snakebites and scorpion stings in Africa: a crucial step for the management of envenomation. Journal of Venomous Animals and Toxins including Tropical Diseases (2016) 22:11. doi: 10.1186/s40409-016-0062-y.

11. Simpson ID, Norris RL (2009) The global snakebite crisis - a public health issue misunderstood, not neglected. Wilderness and Environmental Medicine 20: 43–56.

12. Chippaux J-P (2017) Snakebite envenomation turns again into a neglected tropical disease! Journal of Venomous Animals and Toxins including Tropical Diseases (2017) 23:38 doi:10.1186/s40409-017-0127-6.

13. Williams D, Gutiérrez JM, Harrison R, Warrell DA, White J, Winkel KD, Gopalakrishnakone P (2010) The global snake bite initiative: an antidote for snake bite. On behalf of the Global Snake Bite Initiative Working Group and International Society on Toxinology. Viewpoint. The Lancet 375: 89–91.

14. Molesworth AM, Harrison R, Theakston RDG, Lalloo DG (2003) Geographic Information System mapping of snakebite incidence in northern Ghana and Nigeria using environmental indicators: a preliminary study. Trans R Soc Trop Med & Hyg 97: 188–192.

15. Hamza M, Idris MA, Maiyaki MB, Lamorde M, Chippaux J-P, Warrell DA, Kuznik A, Habib AG (2016) Cost-effectiveness of antivenoms for snakebite envenoming in 16 countries in West Africa. PLoS Negl Trop Dis 10(3): e0004568. doi:10.1371/journal. pntd.0004568.

16. Lam A, Camara B, Kane O, Diouf A, Chippaux J-P Epidemiology of snakebites in Kédougou region (eastern Senegal): comparison of various methods for assessment of incidence and mortality. Journal of Venomous Animals and Toxins including Tropical Diseases (2016) 22:9. DOI: 10.1186/s40409-016-0064-9.

17. Ghana Statistical Service (2014) - 2010 Population and Housing Census, Analytical Report for the Savelugu-Nanton District. (http://www.statsghana.gov.gh/docfiles/2010_District_Report/Northern/SAVELUGU.pdf).

18. Nkrumah, F, Klutse NAB, Adukpo DC, Owusu K, Quagraine KA, Owusu A, Gutowski Jr W (2014) Rainfall variability over Ghana: Model versus rain gauge observation. International Journal of Geosciences 5: 673–683. (http://dx.doi.org/10.4236/ijg.2014.57060).

19. Ntume R, Anywar G (2015) Ethnopharmacological survey of medicinal plants used in the treatment of snakebites in Central Uganda. Current Life Sciences (2015); 1(1): 6–14.

20. LeCompte MD, Goetz JP (1982) Problems of reliability and validity in ethnographic research. Review of Educational Research 52(1): 31–60. https://doi.org/10.3102/00346543052001031.

21. Jones JPG, Andriamarovololona MM, Hockley N, Gibbons JM, Milner-Gulland EJ (2008) Testing the use of interviews as a tool for monitoring trends in the harvesting of wild species. Journal of Applied Ecology 45: 1205–1212.

22. Visser LE, Kyei-Faried S, Belcher DW (2004) Protocol and monitoring to improve snake bite outcomes in rural Ghana. Transactions of the Royal Society of Tropical Medicine and Hygiene 98: 278–283. doi:10.1016/S0035-9203(03)00065-8.

23. Punguyire D, Baiden F, Nyuzaghl J, Hultgren A, Berko Y, Brenner S, Soghoian S, Adjei G, Niyogi A, Moresky R (2014) Presentation, management, and outcome of snake-bite in two district hospitals in Ghana. Pan African Medical Journal 2014; 19:219 doi:10.11604/pamj.2014.19.219.5267.

24. Mensah EK, Karikari K, Aikins M, Vanotoo L, Sackey S, Ohuabunwo C, Wurapa F, Sifah TK, Afari E Secondary analysis of snake bite data in the Western Region of Ghana: 2006-2010. Ghana Med J 2016; 50(2): 103–106 DOI: http://dx.doi.org/10.4314/gmj.v50i2.8.

25. Habib AG, Abubakar SB (2011) Factors affecting snakebite mortality in north-eastern Nigeria. International Health 3:50–55.

26. Tochie JN, Temgoua MN, Njim T, Celestin D, Tankeu R, Nkemngu NJ The neglected burden of snakebites in Cameroon: a review of the epidemiology, management and public health challenges. BMC Res Notes (2017) 10:405.

27. Gampini S, Nassouri S, Chippaux J-P, Semde Rasmané (2016) Retrospective study on the incidence of envenomation and accessibility to antivenom in Burkina Faso. Journal of Venomous Animals and Toxins including Tropical Diseases (2016) 22:10. DOI: 10.1186/s40409-016-0066-7.

28. Tagwireyi DB, Ball DE, Nhachi CFB (2006) Differences and similarities in poisoning admissions between urban and rural health centers in Zimbabwe. Clinical Toxicology 44: 233–241.

29. Mbarouk GS, Sawe HR, Mfinanga JA, Stein J, Levin S, Mwafongo V, Runyon MS, Reynolds TA, Olson KR. Patients with acute poisoning presenting to an urban emergency department of a tertiary hospital in Tanzania. BMC Res Notes (2017) 10:482. DOI: 10.1186/s13104-017-2807-2.

30. Chippaux J-P Estimate of the burden of snakebites in sub-Saharan Africa: A meta-analytic approach. Toxicon 57 (2011) 586–599.

31. Sani UM, Jiya NM, Ibitoye PK, Ahmad MM (2013) Presentation and outcome of snake bite among children in Sokoto, North-Western Nigeria. Sahel Medical Journal 16:148–53.

32. Ghana Health Service (2017) The Health Sector in Ghana: Facts and Figures. Centre for Health and Information Management (CHIM), Policy Planning, Monitoring and Evaluation Division (PPMED). Report by collaborative partners; Ghana Health Service, Ministry of Health, Ghana Statistical Services, World Health Organization. 60 pp. https://www.ghanahealthservice.org/downloads/FACTS+FIGURES_2017.pdf.

33. Schioldann E, Mahmood MA, Kyaw MM, Halliday D, Thwin KT, Chit NN, et al. (2018) Why snakebite patients in Myanmar seek traditional healers despite availability of biomedical care at hospitals? Community perspectives on reasons. PLoS Negl Trop Dis 12(2): e0006299. https://doi.org/10.1371/journal.pntd.0006299.

34. Sloan DJ, Dedicoat MJ, Lalloo DG (2007) Health-Seeking behaviour and use of traditional healers after snakebite in Hlabisa sub-district, KwaZulu Natal. Tropical Medicine and International Health 12(11): 1386–1390.

35. Habib AG (2013) Public health aspects of snakebite care in West Africa: perspectives from Nigeria. Journal of Venomous Animals and Toxins including Tropical Diseases 19:27. http://www.jvat.org/content/19/1/27.

36. Molander M, Staerk D, Nielsen HM, Brandner JM, Diallo D, Zacharie CK, van Staden J, Jäger AK (2015) Investigation of skin permeation, ex vivo inhibition of venom-induced tissue destruction, and wound healing of African plants used against snakebites. Journal of Ethnopharmacology 165:1–8. http://dx.doi.org/10.1016/j.jep.2015.02.014.

37. Molander M, Saslis-Lagoudakis CH, Jäger AK, Rønsted N (2012) Cross-cultural comparison of medicinal floras used against snakebites. Journal of Ethnopharmacology 139:863–872. doi:10.1016/j.jep.2011.12.032.

38. Chekole G (2017) Ethnobotanical study of medicinal plants used against human ailments in Gubalafto District, Northern Ethiopia. Journal of Ethnobiology and Ethnomedicine (2017) 13:55 DOI 10.1186/s13002-017-0182-7.

39. Nnamani, CV, Ukwa EV (2015) Taxonomic diversity of medicinal plants utilized for traditional management of snakebite in southeast, Nigeria: Conservation for sustainability. International Journal of Development and Sustainability 4(12):1138–1152.

40. Habib AG, Kuznik A, Hamza M, Abdullahi MI, Chedi BA, Chippaux J-P, et al. (2015) Snakebite is Under Appreciated: Appraisal of Burden from West Africa. PLoS Negl Trop Dis 9(9): e0004088. doi:10.1371/journal.pntd.0004088.

41. Brown NI (2012) Consequences of Neglect: Analysis of the Sub-Saharan African Snake Antivenom Market and the Global Context. PLoS Negl Trop Dis 6(6): e1670. doi:10.1371/journal.pntd.0001670.

42. Visser LE, Kyei-Faried S, Belcher DW, Geelhoed DW, van Leeuwen JS, van Roosmalen J (2008) Failure of a new antivenom to treat Echis ocellatus snake bite in rural Ghana: the importance of quality surveillance. Transactions of the Royal Society of Tropical Medicine and Hygiene 102:445–450. doi:10.1016/j.trstmh.2007.11.006.

43. Habib AG, Gebi UI, Onyemelukwe GC (2001) Snake bite in Nigeria. Afr J Med Med Sci. 30(3):171–8.

44. Omogbai, EKI, Nworgu ZAM, Imhafidon MA, Ikpeme AA, Ojo DO, Nwako CN (2002) Snakebites in Nigeria: A study of Prevalence and Treatment in Benin City. Tropical Journal of Pharmaceutical Research:1(1): 39–44.

45. Ogunfowokan O (2012) Bite-to-hospital time and morbidity in victims of Viper bite in a rural hospital in Nigeria. African Journal Primary Health Care & Family Medicine 4(1), Art. #371, 7 pages. http://dx.doi.org/10.4102/phcfm.v4i1.371.

46. Paramonte B (2007) Snake bites in Nigeria. Medical Journal of Therapeutics Africa. 1(3):222–226.

47. Fadare JO, Afolabi OA (2012) Management of snake bite in resource-challenged setting: A review of 18 months experience in a Nigerian hospital. Journal of Clinical Medicine and Research 4(3):39–43. http://www.academicjournals.org/JCMR

48. Chafiq F, Hattimy FE, Rhalem N, Chippaux J-P, Soulaymani A, Mokhtari A, Soulaymani-Bencheikh R (2016) Snakebites notified to the poison control center of Morocco between 2009 and 2013 Journal of Venomous Animals and Toxins including Tropical Diseases 22:8. DOI 10.1186/s40409-016-0065-8.

49. Hattimy FE, Chafiq F, Hami H, Mokhtari A, Soulaymani A, Soulaymani-Bencheikh R (2018) Geographical distribution of health indicators related to snake bites and envenomation in Morocco between 1999 and 2013. Epidemiology and Health 40:e2018024. https://doi.org/10.4178/epih.e2018024.

50. Ochola FO, Okumu MO, Muchemi GM, Mbaria JM, Gikunju JK (2018) Epidemiology of snake bites in selected areas of Kenya. Pan African Medical Journal 29:217. doi:10.11604/pamj.2018.29.217.15366.

51. Sakajiki A, Ilah G, Lukman A-A, Yakasai A (2017) Snake bite envenomation seen at a specialist hospital in Zamfara state, North-Western Nigeria. Annals of Tropical Medicine and Public Health: 10(2)p.391. Academic One File.

52. Nkwescheu A, Mbasso LCD, Pouth FBB, Dzudie A, Billong SC, Ngouakam H, Diffo JLD, Eyongorock H, Mbacham W (2016) Snakebite in bedroom kills a physician in Cameroon: a case report. Pan African Medical Journal 24:231. doi:10.11604/pamj.2016.24.231.7576. http://www.panafrican-med-journal.com/content/article/24/231/full/.

53. Tianyi F-L, Agbor VN, Tochie JN, Kadia BM, Nkwescheu AS (2018) Community-based audits of snake envenomations in a resource-challenged setting of Cameroon: case series. BMC Res Notes 11:317. https://doi.org/10.1186/s13104-018-3409-3.

54. Tianyi F-L, Dimala CA, Feteh VF (2017) Shortcomings in snake bite management in rural Cameroon: a case report. BMC Res Notes 10:196. DOI 10.1186/s13104-017-2518-8.

55. Enwere, GC, Obu HA, Jobarteh A (2000) Snake bites in children in The Gambia. Annals of Tropical Paediatrics. 20(2):121–124. ISSN 0272-4936 print/ISSN 1465-328/online/00/020121-04.

56. Ruiz-Casares M (2009) Unintentional Childhood Injuries in Sub-Saharan Africa: An Overview of Risk and Protective Factors. Journal of Health Care for the Poor and Underserved 20(4) Suppl.:51–67 (Article). Johns Hopkins University Press. DOI: https://doi.org/10.1353/hpu.0.0226.

57. Butler JA (1993) Seasonal Reproduction in the African Olive Grass Snake, Psammophis phillipsi (Serpentes: Colubridae). Journal of Herpetology 27(2):144–148.

58. Akani GC, Eniang EA, Ekpo IJ, Angelici FM, Luiselli L (2003) Food Habits of the Snake Psammophis phillipsi from the Continuous Rain-Forest Region of Southern Nigeria (West Africa). Journal of Herpetology 37(1):208–211.

59. Yahaya M (2010) The occurrence, distribution, diversity and toxicology of snakes in the Savelugu-Nanton District of the Northern Region of Ghana. MPhil. Thesis. Department of Animal Biology and Conservation Science, University of Ghana, Legon, Accra. 147 pages.

